# Spatiotemporal kinetics of CAF-1-dependent chromatin maturation ensures transcription fidelity during S-phase

**DOI:** 10.1101/2023.05.25.541209

**Authors:** Boning Chen, Heather K. MacAlpine, Alexander J. Hartemink, David M. MacAlpine

## Abstract

Proper maintenance of epigenetic information after replication is dependent on the rapid assembly and maturation of chromatin. Chromatin Assembly Complex 1 (CAF-1) is a conserved histone chaperone that deposits (H3-H4)_2_ tetramers as part of the replication-dependent chromatin assembly process. Loss of CAF-1 leads to a delay in chromatin maturation, albeit with minimal impact on steady-state chromatin structure. However, the mechanisms by which CAF-1 mediates the deposition of (H3-H4)_2_ tetramers and the phenotypic consequences of CAF-1-associated assembly defects are not well understood. We used nascent chromatin occupancy profiling to track the spatiotemporal kinetics of chromatin maturation in both wild-type (WT) and CAF-1 mutant yeast cells. Our results show that loss of CAF-1 leads to a heterogeneous rate of nucleosome assembly, with some nucleosomes maturing at near WT kinetics and others exhibiting significantly slower maturation kinetics. The slow-to-mature nucleosomes are enriched in intergenic and poorly transcribed regions, suggesting that transcription-dependent assembly mechanisms can reset the slow-to-mature nucleosomes following replication. Nucleosomes with slow maturation kinetics are also associated with poly(dA:dT) sequences, which implies that CAF-1 deposits histones in a manner that counteracts resistance from the inflexible DNA sequence, promoting the formation of histone octamers as well as ordered nucleosome arrays. In addition, we demonstrate that the delay in chromatin maturation is accompanied by a transient and S-phase specific loss of gene silencing and transcriptional regulation, revealing that the DNA replication program can directly shape the chromatin landscape and modulate gene expression through the process of chromatin maturation.

## Introduction

In eukaryotic cells, genomic DNA is packaged into chromatin, a highly organized DNA-protein complex that encodes the epigenetic information essential for regulating DNA-mediated processes, including gene transcription (Jiang and Pugh 2009), DNA replication (Kurat et al. 2017) and DNA repair (Chen and Tyler 2022). Despite the critical regulatory role of chromatin, the epigenetic landscape is disrupted every cell cycle during S-phase. Specifically, as replication forks proceed through the genome, parental chromatin ahead of the fork must be disassembled for DNA synthesis and then re-established in the wake of the fork (Jackson 1990; Gruss et al. 1993). Efficient and faithful propagation of the chromatin structure is critical for maintaining genome integrity and epigenetic memory.

Genetic and biochemical experiments have identified many factors and mechanisms involved in replication-coupled chromatin assembly (MacAlpine and Almouzni 2013). The assembly of nucleosomes is a step-wise process starting with the deposition of (H3-H4)_2_ tetramers, either recycled from parent chromatin or newly synthesized, followed by the addition of two H2A-H2B dimers (Smith and Stillman 1991; Jackson 1988). Nascent and parental (H3-H4)_2_ tetramers carry distinct histone post-translational modifications (PTM) and are deposited onto the leading and lagging strand in a coordinated manner that ensures proper inheritance of the PTM landscape in the daughter cells (Masumoto et al. 2005; Reverón-Gómez et al. 2018; Li et al. 2020; Gan et al. 2018). Under certain conditions, (H3-H4)_2_ tetramers can form stable intermediates with DNA termed tetrasomes (Hall et al. 2009; Andrews et al. 2010; Böhm et al. 2011; Ordu et al. 2019), but typically the formation of intact nucleosomes occurs rapidly and is tightly coupled to the replication fork (Gasser et al. 1996; Cusick et al. 1984; Sogo et al. 1986).

Chromatin assembly is predominantly mediated by histone chaperones, and one of the key players is CAF-1 (Smith and Stillman 1991; Gurard-Levin et al. 2014; Miller and Costa 2017). CAF-1 is a three-subunit protein complex consisting of Cac1, Cac2, and Cac3 (p150, p60, and p48 in human cells) and is highly conserved from yeast to human cells. Its role in promoting nucleosome assembly on nascent DNA strands was first identified in an SV40-based *in vitro* study (Smith and Stillman 1989). More recent studies found that CAF-1 receives nascent H3-H4 dimers from Asf1, facilitates the dimerization of H3-H4 to form (H3-H4)_2_ tetramers and deposits them onto the newly replicated DNA through an interaction with PCNA (Shibahara and Stillman 1999; Liu et al. 2016, 2017). In addition, CAF-1 exhibits a high affinity for the post-translational modification H3K56ac, present on nascent H3-H4 in yeast, suggesting that CAF-1 contributes to the deposition of new histones (Li et al. 2008; Masumoto et al. 2005).

CAF-1 is essential in muti-cellular organisms. Loss of CAF-1 causes development to cease at the early embryonic stage in mice and leads to larval lethality in *Drosophila (Houlard et al. 2006; Yu et al. 2013).* In human cells, loss of CAF-1 halts S-phase progression and arrests cells in early or mid S-phase (Hoek and Stillman 2003). In yeast, disruption of CAF-1 does not affect the viability of the cells, which could be due, in part, to the functional redundancy of proteins commonly found in yeast. However,

CAF-1 mutant yeast cells do display phenotypic defects, including loss of silencing at the subtelomeric regions and mating type loci, as well as increased sensitivity to UV damage (Kaufman et al. 1997; Enomoto and Berman 1998; Monson et al. 1997; Smith et al. 1999). Despite the central role of CAF-1 in chromatin assembly, loss of CAF-1 in yeast has minimal impact on nucleosome organization and positioning (van Bakel et al. 2013; Fennessy and Owen-Hughes 2016). While the molecular function of CAF-1 is well-established, it remains unclear how loss of CAF-1-associated phenotypes relates to its role in chromatin assembly.

To better understand the role of CAF-1 in chromatin maturation, we monitored the spatiotemporal kinetics of chromatin maturation in WT and CAF-1 mutant cells. We used nucleotide analog labeling and chromatin occupancy profiling to monitor the kinetics of chromatin maturation throughout the genome (Fennessy and Owen-Hughes 2016; Vasseur et al. 2016; Ramachandran and Henikoff 2016; Stewart-Morgan et al. 2019; Gutiérrez et al. 2019). Our results show that CAF-1 is essential for timely chromatin maturation, as loss of CAF-1 causes a pronounced delay in the re-establishment of nucleosome occupancy and organization after replication. Individual nucleosomes were assembled in a heterogeneous fashion, with some being deposited immediately after passing of the replication forks, whereas others were significantly delayed. Despite the significant difference in chromatin maturation kinetics between WT and CAF-1 mutant cells, the steady-state mature chromatin occupancy profiles were nearly indistinguishable. Finally, we demonstrate that rapid and efficient CAF-1-dependent chromatin maturation is essential for establishing epigenetic silencing and suppressing cryptic transcription during S-phase.

## Results

### Nascent chromatin occupancy profiling reveals an assembly defect in cac1Δ cells

To capture the dynamics of chromatin assembly behind the replication fork, we used nascent chromatin occupancy profiling (NCOP) (Gutiérrez et al. 2019). NCOP combines EdU labeling of newly synthesized DNA with MNase profiling (Henikoff et al. 2011) to track the spatiotemporal kinetics of chromatin assembly at nucleotide resolution. An asynchronous population of cells were pulse-labeled with the nucleoside analog, EdU, for 5 minutes, followed by a thymidine chase for 10, 15, 20, or 40 minutes to capture chromatin assembly dynamics (Figure 1A). After the pulse-chase labeling, the chromatin samples were digested by MNase. The EdU-labeled DNA fragments were biotinylated with click chemistry and enriched using streptavidin-biotin isolation before being subjected to paired-end sequencing. DNA fragments protected by the histone octamer should be ∼150 bp in length, whereas small DNA binding factors such as transcription factors or the origin recognition complex protect DNA fragments smaller than 80 bp. Importantly, this assay provides a factor-agnostic view of chromatin assembly dynamics throughout the genome.

**Figure 1.**
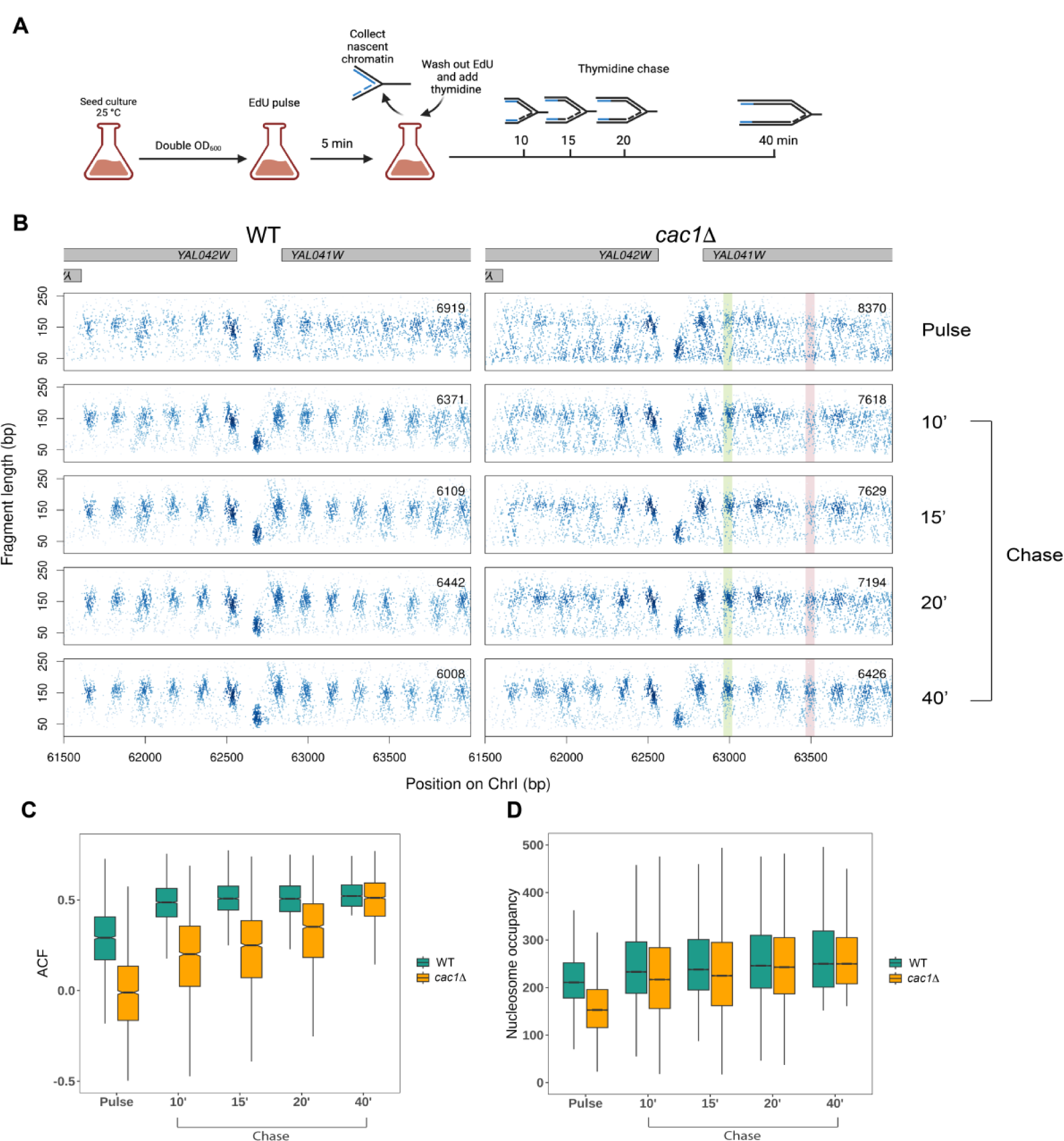
Nascent chromatin occupancy profiling reveals a chromatin maturation defect in *cac1Δ* cells. (A) Schematic of experimental design for capturing chromatin maturation dynamics behind the replication fork. (B) Chromatin occupancy profiles at a representative locus in WT and *cac1Δ* cells after a 5 minute EdU pulse and 10, 15, 20, and 40 minutes following a thymidine chase. Each dot represents a fragment midpoint, the shading of which is determined by a 2D kernel density estimate. The number of fragments in each window is inset on each panel. Gene bodies are shown in gray on the top. An example of a slow-to-mature nucleosome is highlighted in red, and an example of a fast-maturing nucleosome is highlighted in green. (C) Autocorrelation function (ACF) values at each time point for the 2097 genes with regularly phased nucleosome arrays in mature chromatin, defined as genes with ACF greater than the median in the WT 40 minute chase sample. (D) Nucleosome occupancy at all time points for the 41663 high-confidence nucleosomes, defined as nucleosomes with occupancy greater than the 25th percentile in both WT and *cac1Δ* cells following the 40 minute chase period.

Nascent chromatin occupancy profiling of WT cells revealed that EdU-labeled nascent chromatin at the fork, following the short pulse period, was more disorganized relative to the chase series. Specifically, the midpoints of the recovered fragments in the chase series of 10, 15, 20, and 40 minutes were largely localized to a periodic array of focal clusters at ∼150 bp representing nucleosome positions. In contrast, the fragments in the pulse sample were more dispersed relative to the nucleosome positions (Figure 1B). We used the autocorrelation function (ACF) to quantify the transient disorganization of chromatin within gene bodies during the pulse and its recovery through the subsequent chase periods. The autocorrelation function (ACF) reveals the periodicity of a complex signal over time, or in this case, the correlated phasing of nucleosome dyads at an inter-nucleosome distance of 172 bp (Gutiérrez et al. 2019). The chromatin recovered from the pulse period of WT cells has a lower ACF than the subsequent chase periods (Figure 1C; green).

To better understand how CAF-1 affects chromatin maturation dynamics, we also generated NCOPs in cells lacking Cac1, the largest subunit of CAF-1. In budding yeast, the deletion of *Cac1* (*cac1Δ*) is sufficient to abrogate the histone chaperone activity of CAF-1 without affecting the viability of the cells (Liu et al. 2016; Yadav and Whitehouse 2016). The recovered chromatin from the 40 minute chase period in *cac1Δ* cells was nearly indistinguishable from WT chromatin (Figures 1B and 1C). Despite the similarities in mature chromatin, the *cac1Δ* cells exhibited a marked delay in the kinetics of chromatin maturation. The pulse, 10, 15, and 20 minute samples were all significantly less structured than mature chromatin.

We also examined the kinetics of deposition for individual nucleosomes. We identified ∼42,000 high-confidence nucleosomes in WT and *cac1Δ* cells and calculated their occupancy in the two strains throughout the time course (Figure 1D). While the nucleosome occupancy at the final time point is comparable between the two strains, it takes longer for nucleosomes to reach full occupancy in *cac1Δ* cells (Figure 1D), consistent with a defect in the deposition of nascent (H3-H4)_2_ tetramers. Together, these results suggest a transient defect in replication-coupled chromatin assembly and maturation in *cac1Δ* cells.

### Heterogeneous rate of histone octamer assembly behind the fork

In addition to the pronounced delay in chromatin maturation, we also observed that individual nucleosomes in *cac1Δ* cells mature at a heterogenous rate. For example, the nucleosome at 63,000 bp (highlighted in green in Figure 1B) on Chr I is deposited rapidly behind the fork and reaches maximal occupancy 10 minutes after the passing of the replication fork. In contrast, the nucleosome at 63,500 bp (highlighted in red in Figure 1B, Supplemental Figure 1) does not fully mature until 40 minutes after passing of the fork. To systematically quantify the maturation rate for individual nucleosomes, we used the reciprocal of the mean weighted sum of occupancy from each time point to generate an assembly timing index (ATI), with lower values reflecting slower maturation kinetics (Van Rechem et al. 2021; Raghuraman et al. 2001). We calculated an ATI value for all high-confidence nucleosomes in *cac1Δ* and WT cells, respectively, and plotted their distributions (Figure 2B). The lower ATI from *cac1Δ* cells is consistent with a global delay in nucleosome deposition. To visualize the maturation dynamics of individual nucleosomes, we generated ordered heatmaps, based on the ATI value, depicting the nucleosome occupancy at each time point relative to the final time point (Figure 2A). For visualization purposes, we are only depicting the nucleosomes from ChrIV, but they are representative of the nucleosomes throughout the genome (Supplemental Figure 2). The heatmaps for WT and *cac1Δ* were ordered independently, as we found no correlation in ATI values between individual nucleosomes (Supplemental Figure 3). In contrast to the uniform maturation pattern seen in WT cells, *cac1Δ* cells exhibit diverse maturation dynamics.

**Figure 2.**
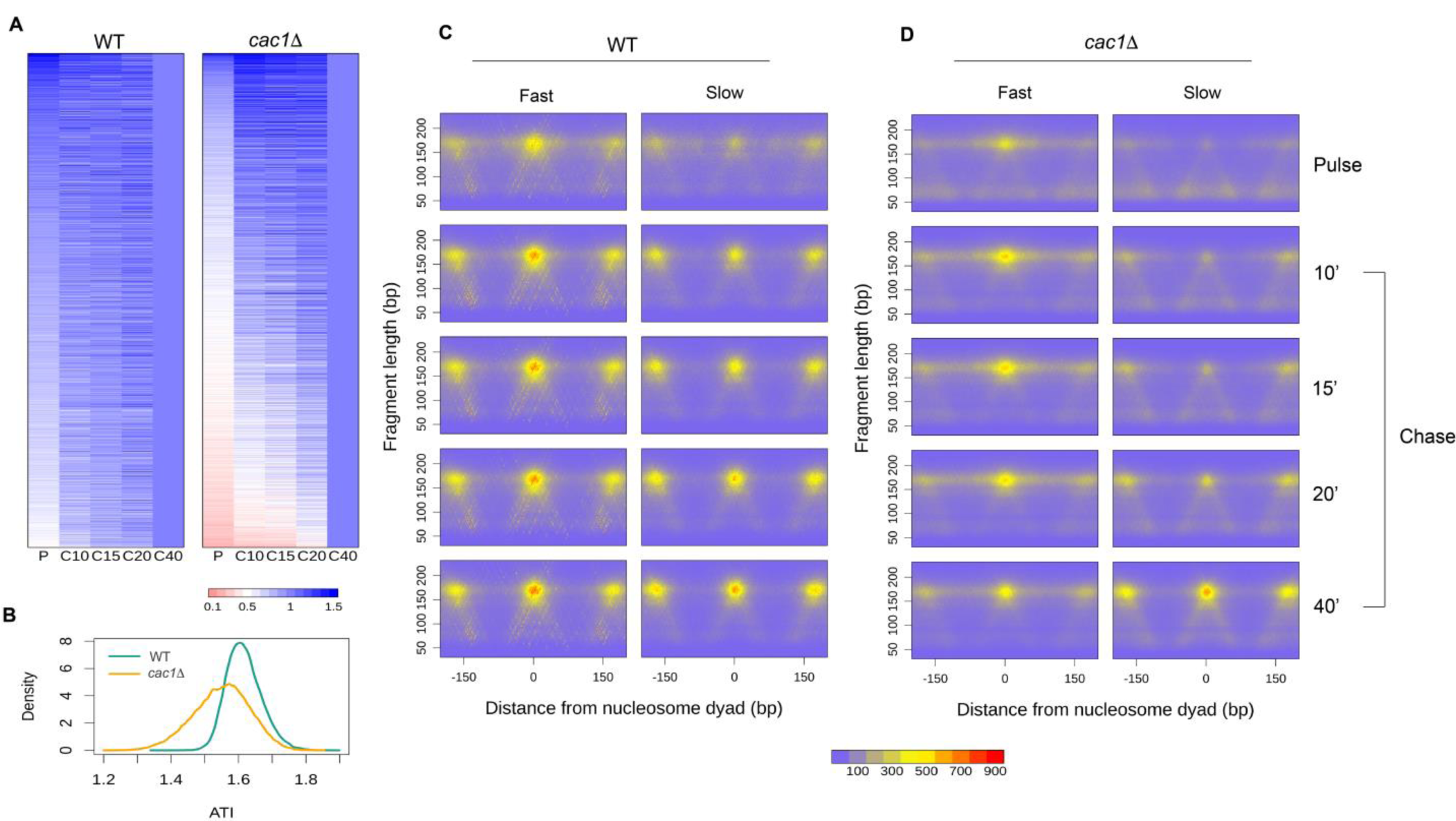
*cac1Δ* cells exhibit a heterogeneous rate of histone octamer assembly behind the fork. (A) Heat maps of nucleosome occupancy at each time point relative to the final time point for the 5344 high-confidence nucleosomes on ChrIV. Each row represents an individual nucleosome. Heat maps are ordered independently by decreasing ATI values. (B) Density distributions of ATI values for all high-confidence nucleosomes in WT and *cac1Δ* cells. (C, D) Aggregated chromatin profiles at all time points for the fast and slow nucleosomes identified in (C) WT and (D) *cac1Δ* cells.

To further explore the differences in maturation dynamics, we sought to agnostically identify the slowest and fastest maturing nucleosomes. We first divided the nucleosomes into deciles based on occupancy in mature chromatin to ensure that fast and slow nucleosomes were comparable based on their final occupancy level. From each decile, we defined nucleosomes with an ATI below the 10th percentile as slow nucleosomes and those with an ATI above the 90th percentile as fast nucleosomes. We identified 4163 fast and slow nucleosomes from both WT and *cac1Δ* cells. We generated aggregate chromatin occupancy profiles from the slow and fast nucleosomes by aligning to the nucleosome dyads of either the fast or slow nucleosomes (Figure 2C). As expected, in WT cells, the occupancy profiles from slow and fast nucleosomes were highly similar throughout the time course. The lack of distinction between slow and fast nucleosomes in WT is consistent with the homogenous maturation pattern we observed. In contrast, in *cac1Δ* cells, the occupancy profiles of slow and fast nucleosomes were significantly different, depicting two populations of nucleosomes with distinct maturation kinetics (Figure 2D, Supplemental Figure 4). The occupancy of fast nucleosomes plateaus within 10 minutes, whereas the slow nucleosomes mature more gradually over the time course.

Several groups have reported the existence of ‘fragile nucleosomes’ in yeast that are more sensitive to MNase digestion (Knight et al. 2014; Henikoff et al. 2011; Kubik et al. 2015; Weiner et al. 2010; Xi et al. 2011; Chereji et al. 2017). To confirm that differences in maturation rate observed in *cac1Δ* cells were not due to fragile nucleosomes and increased sensitivity to MNase digestion, we calculated the ATI values for previously identified fragile nucleosomes (Mitra et al. 2021; Chereji et al. 2017) in both WT and *cac1Δ* cells (Supplemental Figure 5). The density distribution of ATI values for fragile nucleosomes is similar to the full set of high-confidence nucleosomes in both WT and *cac1Δ* cells. We saw no enrichment of fragile nucleosomes that may represent slow nucleosomes in either WT or *cac1Δ* cells, suggesting that the nucleosome assembly rate is independent of its sensitivity to MNase digestion. Overall, we conclude that *cac1Δ* cells exhibit a heterogeneous rate of nucleosome maturation behind the fork as a result of the chromatin assembly defect.

### Genomic and sequence features associated with slow-maturing nucleosomes

To explore the genomic and sequence features influencing the rate of nucleosome deposition in *cac1Δ,* we examined the distribution of slow nucleosomes throughout the yeast genome. The distribution of slow-maturing nucleosomes along each of the chromosomes appeared random, with a median inter-slow nucleosome distance of 1050 bp; however, we did note a bias, or enrichment, of slow nucleosomes in intergenic regions relative to gene bodies (p < 2.2 x 10^-16^, Supplemental Figure 6). Chromatin is assembled by both replication-dependent and replication-independent mechanisms. We hypothesized that transcription-dependent chromatin assembly mechanisms (Ray-Gallet et al. 2002) may reset the gene body chromatin, especially for transcripts that are highly expressed, including during S-phase (Vasseur et al. 2016; Stewart-Morgan et al. 2019).

To explore the relationship between the density of slow nucleosomes within a gene and gene expression, we binned the genes by quintiles of gene expression and generated boxplots depicting the density of slow nucleosomes within each gene (Figure 3A). We found that the least expressed genes (quintile 1) exhibited a slow nucleosome density that was similar to intergenic sequences suggesting a default pattern of deposition behind the replication fork. In contrast, with increasing transcription, we observed a gradual decrease in the density of slow nucleosomes within gene bodies. Together, these results suggest that the heterogeneous patterns of nucleosome deposition we observe in the absence of Cac1 are dependent on replication and that active transcription and transcription-dependent chromatin assembly mechanisms can rapidly restore the chromatin to its native and mature state.

**Figure 3.**
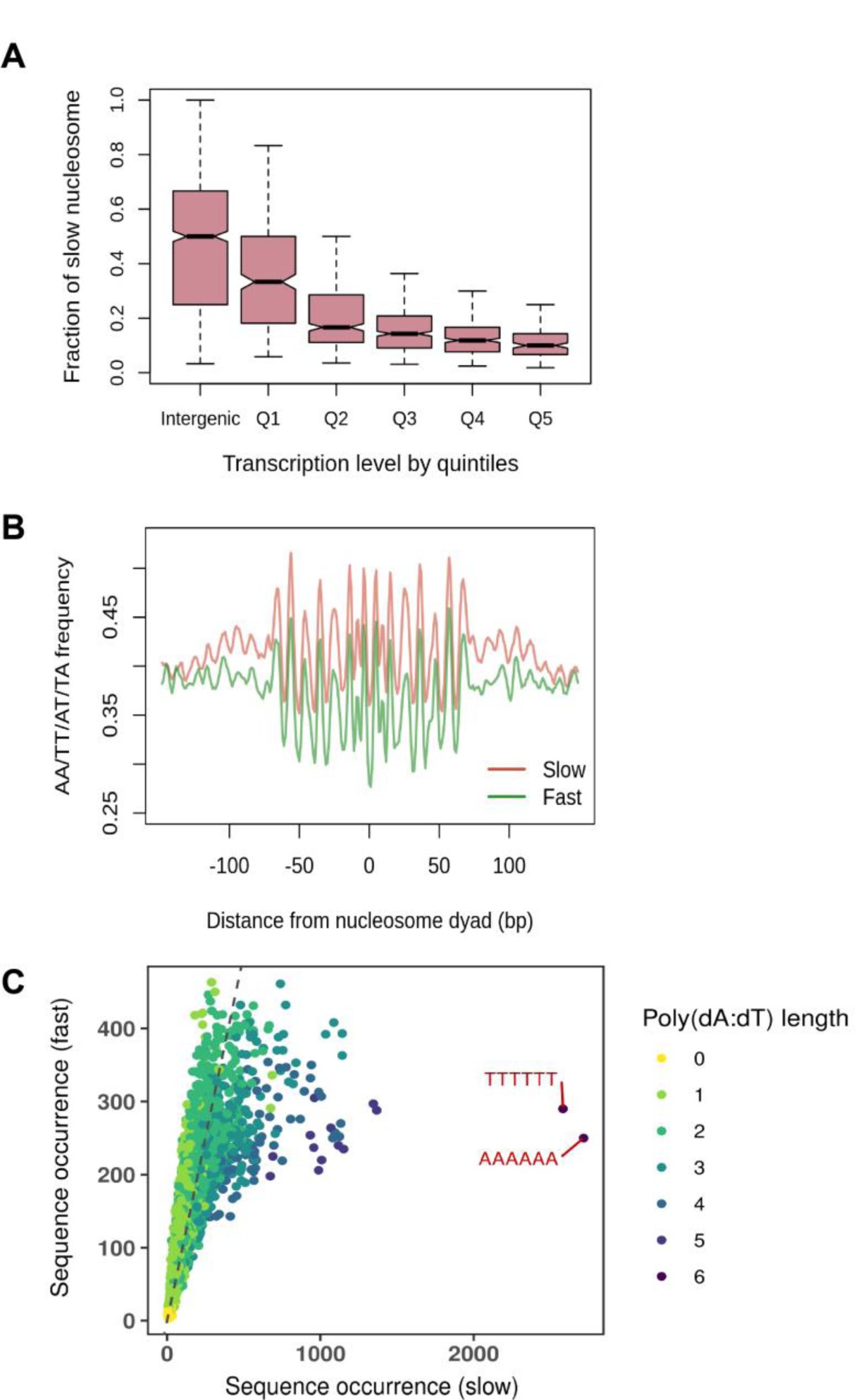
Genomic and sequence features associated with slow-to-mature nucleosomes. (A) Fraction of slow nucleosomes identified in *cac1Δ* cells among all nucleosomes in the intergenic regions and gene bodies. Genes are binned into quintiles based on expression levels <u>(Churchman and Weissman 2011)</u>. Q1 represents genes with the lowest expression level, and Q5 represents genes with the highest expression level. (B) Aggregated position-dependent A/T dinucleotide frequencies surrounding the dyads of fast and slow nucleosomes. (C) Scatterplot depicting the occurrence of all possible 6-mer sequences from the DNA associated with fast and slow nucleosomes. Dots are colored by the length of the longest poly(dA:dT) sequence that exists in the sequence. The dotted grey line denotes the diagonal line.

Aside from the rapid transcription-dependent re-establishment of chromatin at the most actively transcribed genes, we did not observe a discernable pattern in the localization of slow-to-mature nucleosomes; thus, we reasoned that other more localized features might govern the maturation speed of individual nucleosomes. It is well established that histone octamers have a strong sequence preference, largely due to the bendability of discrete nucleotide sequences, which is crucial for the formation of nucleosomes (Drew and Travers 1985). Optimal nucleosome formation occurs when the more bendable A/T dinucleotides exist in a ∼10 bp periodicity (Segal et al. 2006; Ioshikhes et al. 2011). In contrast, poly(dA:dT) sequences are inherently stiff, impeding nucleosome formation (Segal and Widom 2009; Nelson et al. 1987). To determine whether local sequence context influences the maturation dynamics of nucleosomes, we determined whether slow and fast nucleosomes are enriched for different sequence motifs. Looking at the nucleotide frequency of the sequences covered by the fast and slow nucleosomes, we discovered that fast and slow nucleosomes both display the classic A/T dinucleotide periodicity; however, slow nucleosomes have elevated levels of A/T frequency, suggesting a potential enrichment in poly(dA:dT) sequences (Figure 3B). Indeed, we found that slow nucleosomes feature multiple poly(dA:dT) sequence elements that hinder nucleosome formation (Figure 3C). These results suggest that the histone chaperone function of CAF-1 overcomes the resistance to nucleosome formation from local sequences and ensures a rapid and efficient assembly of nucleosomes at diverse sequences across the genome.

### Sub-nucleosomal fragments captured in *cac1**Δ*** cells reveal nucleosome intermediates

In addition to the perturbed nucleosomal chromatin, the nascent chromatin of *cac1Δ* cells also displayed a genome-wide increase in small fragments that diminished as the chromatin matured (Figure 4A, 4B, Supplemental Figure 7). The small fragments appeared to be present in an array of repeating clusters. Using the autocorrelation function, we found that they exhibited a periodicity of 70 bp (Supplemental Figure 8). Due to the ubiquity and periodicity of the small fragments, we hypothesize that they reflect either loosely bound fragile nucleosomes or nucleosome intermediates (e.g., tetrasomes) and not sites of promiscuous transcription factor binding. The length of the small protected fragments is ∼60 bp which is consistent with the length of DNA predicted to be protected by tetrasomes (Rychkov et al. 2017).

**Figure 4.**
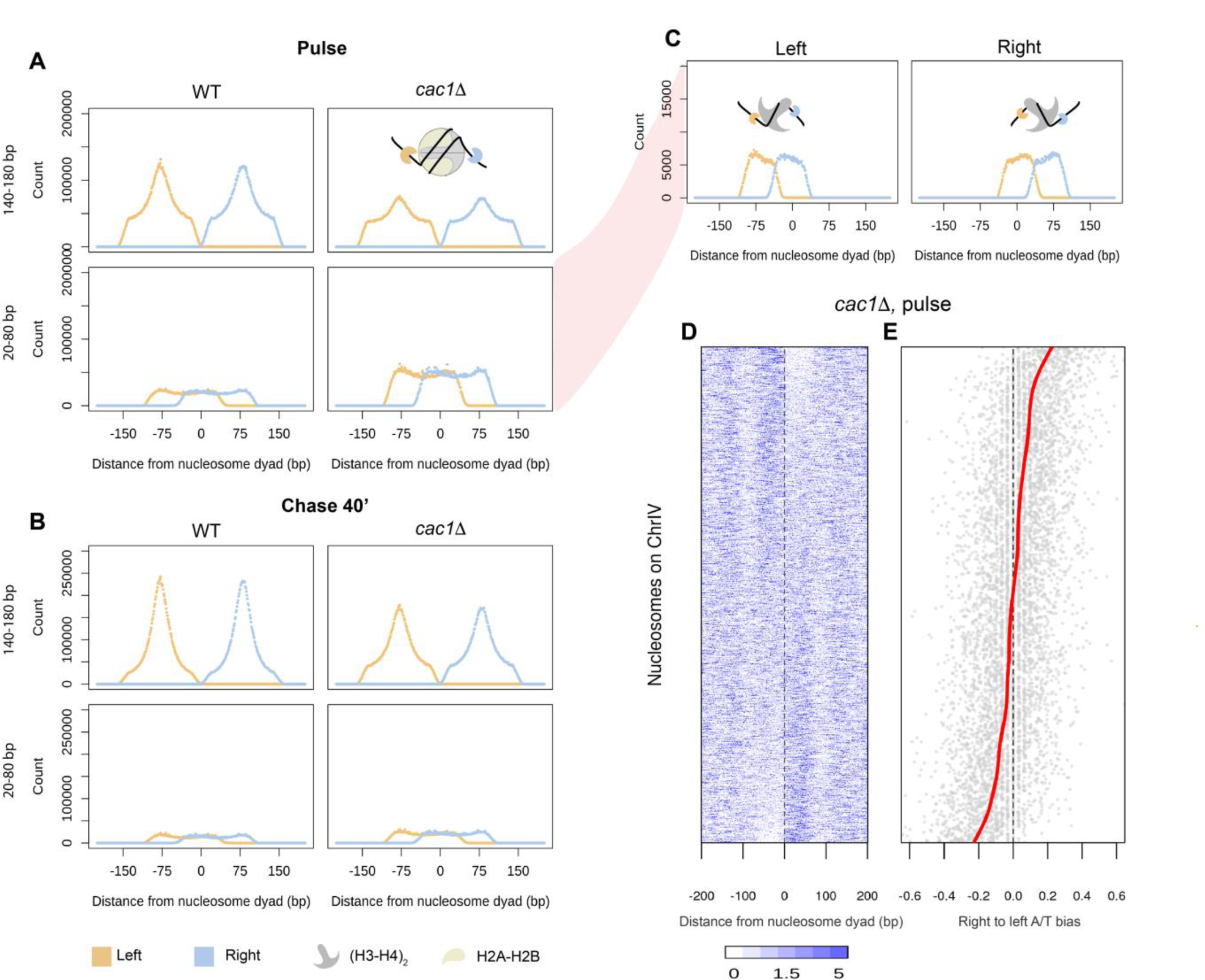
Sub-nucleosomal fragments captured in cac1Δ cells reveal nucleosome assembly intermediates. (A, B) Aggregate signals of the left (orange) and right (blue) ends of fragments surrounding the dyads of high-confidence nucleosomes in (A) nascent chromatin and (B) mature chromatin in the two strains. Fragments are divided into nucleosome-sized fragments (140 - 180 bp in length, top panels) and subnucleosomal-sized fragments (20 - 80 bp in length, bottom panels). A diagram depicting the fragment ends in relation to the nucleosome is inset in the panel. (C) Fragment ends of the subnucleosomal fragments in the nascent chromatin of *cac1Δ* cells are plotted separately by the location of the fragment midpoint relative to the nucleosome dyads. (D) Heatmap depicting occupancy of subnucleosomal fragments surrounding individual nucleosomes. Each row corresponds to a high-confidence nucleosome on ChrIV. Rows are ordered by increasing right-to-left occupancy bias of the subnucleosomal fragments. (E) Dot plot showing the right-to-left A/T bias of DNA sequence associated with individual nucleosomes in the same order as (D). The red line denotes the fitted smooth spline curve.

To explore the hypothesis that these small fragments may represent nucleosome assembly intermediates, we sought to investigate the positioning of the recovered fragments relative to the position of mature nucleosome dyads. To delineate the protected fragments, we visualized the left (orange) and right (blue) ends of each fragment to identify the precise boundaries of the protein-DNA interaction. For example, following the 40 minute chase periods, the ends of nucleosome-sized fragments (140-180 bp) in both WT and *cac1Δ* cells gave rise to two peaks of signal that clearly define the nucleosome edges (Figure 4B). In contrast, during the short 5 min pulse period, the left and right peaks delineating the nucleosome boundary were less defined, especially in *cac1Δ* cells (Figure 4A). Examination of the left and right fragment edges derived from smaller reads (20-80 bp) exhibited a bimodal population with one set of fragments starting at the left edge of the nucleosome and ending at the dyad and the next set of fragments starting at the dyad and ending at the right most nucleosome boundary (Figure 4C). This pattern of the bimodal signal was significantly more prominent in the *cac1Δ* cells and likely represented a true nucleosome intermediate, as the pattern was transient and restricted to the earlier pulse and chase periods. The length of the small fragments and their position relative to the dyad are also consistent with fragments recovered from the MNase digestion of tetrasomes formed *in vitro* (Dong and van Holde 1991). Thus, we interpret the broad increase of subnucleosomal fragments as an accumulation of tetrasomes resulting from assembly defects in *cac1Δ* cells.

The aggregate view of tetrasome positions revealed by the analysis of fragment ends may suggest an equal distribution of tetrasomes on the left and right side of the dyad or an asymmetric bias to either the left or right side. We found that the majority of protected and recovered tetrasome fragments exhibited a strong bias to either the right or left side of the mature nucleosome dyad (Figure 4D). Prior *in vitro* work found that differentially orient hexamers can be generated by modulating the orientation of the Widom 601 in a sequence-specific fashion (Levendosky et al. 2016). To understand why tetramers may exhibit a preference for one side of the nucleosome dyad versus the other, we examined the local sequence context, specifically the AT content. We plotted the bias in AT-content on each side of the nucleosome dyad in the same order as the bias in tetrasome occupancy (Figure 4E). We found that tetrasomes have a preference for the less AT-rich side, likely avoiding poly(dA:dT) sequences. Together, these observations suggest that the (H3-H4)_2_ tetramer deposition is skewed by inherent DNA affinity as a result of the CAF-1 defect, which may further impede the subsequent formation of histone octamers.

### Chromatin assembly defects in the absence of CAF-1 result in a transient and S-phase-specific transcriptional dysregulation

CAF-1 defects can lead to multiple transcriptional program changes, including increased cryptic transcription and loss of silencing at the subtelomeric regions and mating type loci (van Bakel et al. 2013; Enomoto and Berman 1998; Monson et al. 1997). Because *Cac1* deletion impacts the kinetics of chromatin assembly during S-phase and not the steady state or mature chromatin occupancy and positioning, we hypothesized that the transcriptional changes found in *cac1Δ* cells are a direct consequence of the delay in chromatin assembly and maturation. To test this hypothesis, we monitored the transcriptional landscape in WT and *cac1Δ* cells throughout the cell cycle. If transcriptional changes are the result of delayed chromatin maturation, they should occur predominantly in S-phase. Cells were synchronized in late G1 with α-factor and then subsequently released into S-phase. Following release into S-phase, we collected samples for RNA expression analysis every 10 minutes for 60 minutes. Following the 60 minute time point, α-factor was added to the culture again to prevent the cells from entering S-phase in the subsequent cell cycle. Cell cycle progression for each time point was monitored by flow cytometry. WT and *cac1Δ* cells progressed through the cell cycle in a similar manner, with cells beginning to enter S-phase at 40 minutes, and by 60 minutes, most cells had entered into G2/M. At 150 minutes after release from the initial α-factor, and following the re-addition of α-factor at 60 minutes, cells were arrested again in G1-phase (Figure 5A).

**Figure 5.**
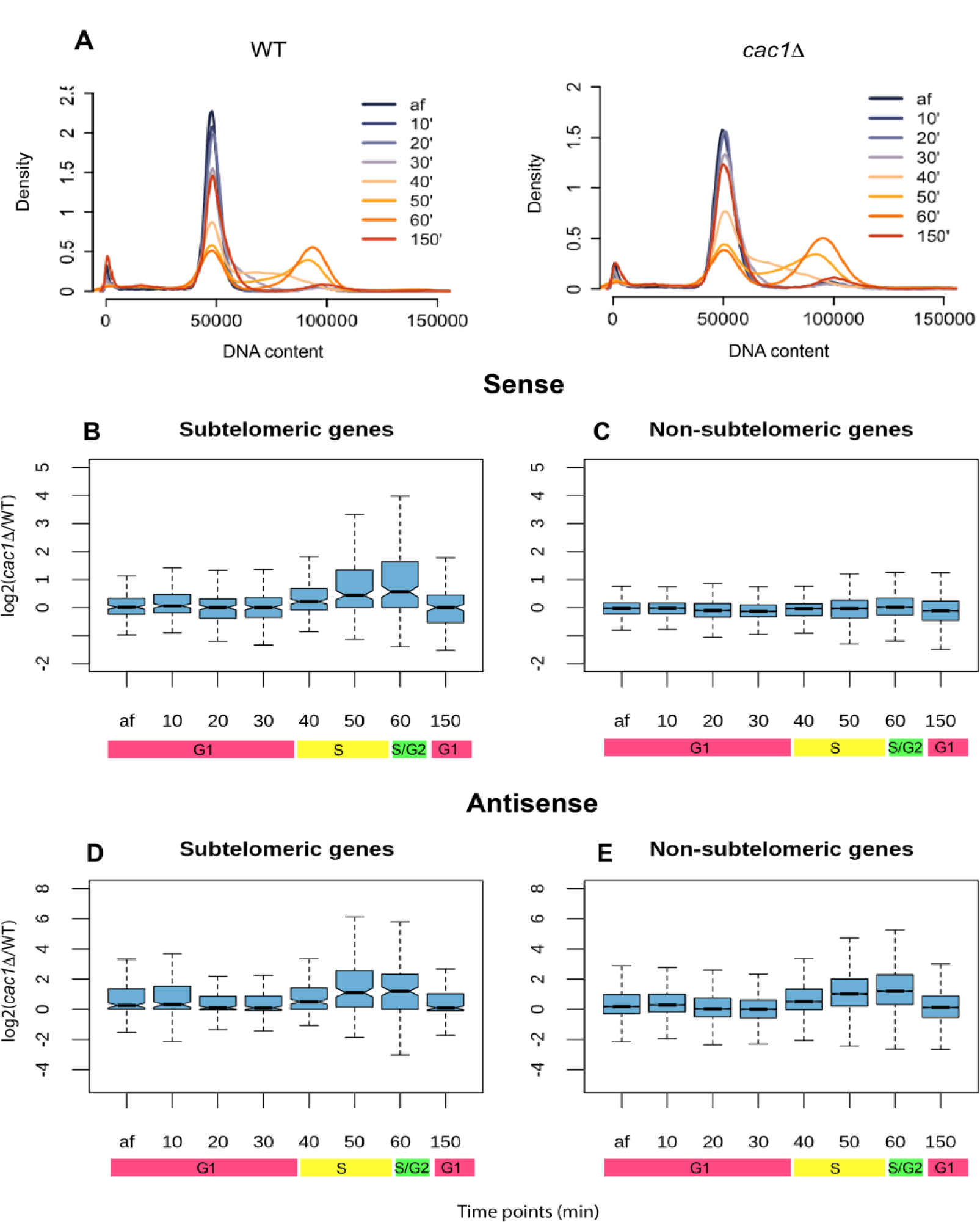
Chromatin assembly defects in the absence of CAF-1 result in a transient and S-phase-specific dysregulation of transcription. (A) DNA content measured by flow cytometry in WT and *cac1Δ* cells at the end of *ɑ*-factor arrest (af) and 10, 20, 30, 40, 50, 60, and 150 minutes after release from the initial *ɑ*-factor arrest. (B-E) Log2 fold-difference of gene expression, measured in TPM, between *cac1Δ* cells and WT cells. (B, C) Sense transcription. (D, E) Antisense transcription. In both cases, genes are divided into subtelomeric genes (B, D) and non-subtelomeric genes (C, E).

We first focused on sense transcription. We observed increased gene expression in *cac1Δ* cells that was specific to subtelomeric genes (located within 20 kb of the chromosome ends) and emerged at 40 minutes, peaked at 60 minutes, and became absent again by 150 minutes, which coincided with the onset and duration of S-phase (Figure 5B, 5C). The increase in subtelomeric gene expression in the absence of CAF-1 activity suggests a CAF1-dependent transient defect in the establishment of subtelomeric silencing. Next, we examined the pattern of antisense transcription throughout the genome. c*ac1Δ* cells exhibited an S-phase-specific increase in antisense transcription. However, unlike sense transcription, the increase in antisense transcription was more widespread and not limited to subtelomeric regions, suggesting different mechanisms for regulating sense and antisense transcription when cells are faced with disruption of the nucleosome organization (Figure 5D, 5E). In summary, our results demonstrate that the elevated antisense transcription and loss of subtelomeric gene silencing occurred in the short window of DNA replication as a result of the delay in chromatin maturation.

## Discussion

The disassembly and ordered reassembly of chromatin at the replication fork are critical for preserving epigenetic memory. We used nascent chromatin occupancy profiling (NCOP) to assess the kinetics of chromatin assembly behind the replication fork in both WT and c*ac1Δ* cells. The spatiotemporal resolution of assembly dynamics throughout the genome revealed a transient defect in the assembly and deposition kinetics of a large fraction of nucleosomes in the absence of the histone chaperone CAF-1. This transient defect in chromatin maturation behind the fork resulted in the dysregulation of transcription during S-phase, including loss of subtelomeric silencing and elevated antisense transcription.

Genetic and biochemical experiments have established the role of CAF-1 in replication-coupled nucleosome assembly (Sauer et al. 2018). However, nucleosome organization and phasing were largely preserved in the absence of CAF-1 subunits (Cac1, Cac2, and Cac3), albeit with a slightly larger linker distance between nucleosomes (van Bakel et al. 2013). More recently, studies in yeast and *Drosophila* have employed an EdU pulse-chase labeling strategy (similar to our NCOP) to map nascent and mature chromatin in the absence of CAF-1. These studies found that mature (chase) chromatin and nucleosome organization, in the absence of CAF-1, were largely indistinguishable from WT chromatin, but that the nascent (pulse) chromatin was more disorganized, suggesting a global defect in chromatin maturation. These studies assessed nucleosome positioning and occupancy in aggregate across the ensemble of gene bodies. While we also observed similar dynamics in aggregate (Supplemental Figure 9), the increased spatial and temporal resolution of our assay allowed us to investigate nucleosome maturation and occupancy on a per-nucleosome basis throughout the genome, which revealed considerable heterogeneity in the assembly kinetics of nucleosomes in the absence of CAF-1. We identified nucleosomes whose maturation rates are near WT (fast nucleosomes), as well as nucleosomes with significantly slower maturation kinetics (slow nucleosomes).

The slow-to-mature nucleosomes occurred throughout the genome with an approximate density of one slow nucleosome per 1 kb. Slow nucleosomes were depleted in gene bodies relative to intergenic sequences. The differential deposition of slow nucleosomes between intergenic regions and gene bodies could be due, in part, to elevated GC content in gene bodies or active transcription. We found a linear relationship between transcriptional activity and the density of slow nucleosomes in gene bodies. Specifically, we found a paucity of slow nucleosomes in actively transcribed genes, while silenced or poorly transcribed genes had a similar density of slow nucleosomes to intergenic regions. These results suggest that the deposition and maturation of slow nucleosomes is a replication-dependent process resulting from the loss of CAF-1 and that transcription-dependent chromatin assembly mechanisms (Ray-Gallet et al. 2002) are able to rapidly reset the chromatin landscape of actively transcribed genes (Vasseur et al. 2016; Stewart-Morgan et al. 2019).

We also explored the sequence features associated with slow nucleosomes and found an enrichment for poly(dA:dT) sequences which are inherently inflexible and prohibitive to the formation of nucleosomes (Segal and Widom 2009; Nelson et al. 1987). We can envision at least two models to account for the presence of slow nucleosomes and their association with poly(dA:dT) sequences. In the first model, CAF-1 activity facilitates the deposition of nascent histone (H3-H4)_2_ tetramers on sequences recalcitrant to nucleosome formation. In the second model, the decreased nucleosome density behind the replication fork resulting from a defect in the deposition of nascent (H3-H4)_2_ tetramers results in the stochastic deposition of histone tetramers on exposed double-stranded DNA without the physical constraints from neighboring nucleosomes. We currently favor the second model as the deposition of parental histones on poly(dA:dT) sequences should not be affected by Cac1 deletion and, therefore, would not result in the strong nucleosome-specific patterns we observe throughout the genome.

We observed an accumulation of regularly phased clusters of small subnucleosomal-sized fragments in the nascent chromatin that was *cac1Δ*-depdendent. The subnucleosomal fragments are approximately 60 bp in length, and the phased clusters of fragments exhibit a periodicity of 70 bp. When aligned to nucleosome dyads, they give rise to two clusters of fragments that lie on either side of the nucleosome dyad, denoting DNA-protein interaction bordered by one edge of the nucleosome and the nucleosome dyad. We interpret the *cac1Δ*-dependent increase in subnucleosomal fragments to suggest that tetrasomes transiently accumulate on nascent chromatin as a result of assembly and maturation defects. Both *in vitro* and *in vivo* experiments demonstrate that nucleosome assembly starts with the binding of (H3-H4)_2_ tetramers to DNA, followed by the binding of H2A-H2B dimers (Wilhelm et al. 1978; Andrews et al. 2010). Considering the geometry of nucleosomes where H2A-H2B dimers surround the central (H3-H4)_2_ tetramers (Luger et al. 1997), the step-wise assembly of nucleosomes implies that the (H3-H4)_2_ tetramers are first deposited at the ‘center’ of the nucleosome, followed by binding of H2A-H2B on either side of the nucleosome. The asymmetric positioning of the tetrasomes captured in *cac1Δ* cells deviates from the proposed step-wise assembly, suggesting that without CAF-1, tetrasomes ‘slide’ to a more thermally stable conformation (indicated by elevated AT-content), which may defer the formation of histone octamers. The positioning of the tetramers is corrected eventually, perhaps by the association of H2A-H2B or the turnover of nucleosomes (Deal et al. 2010; Dion et al. 2007). These results are consistent with the function of CAF-1 in counteracting the inherent DNA-histone affinity preference and anchoring the tetramers at desired positions that promote nucleosome formation. Together, the findings provide important mechanistic insights into understanding how CAF-1 mediates (H3-H4)_2_ tetramer deposition onto nascent DNA.

Following the passage of the DNA replication fork, the parental epigenetic landscape must be restored on both daughter strands. Elegant genetic and biochemical experiments have begun to elucidate factors and mechanisms responsible for the balanced inheritance of parental histones to the leading and lagging strands. Parental histones do not segregate randomly, but rather a diverse collection of factors such as Mcm2, Dpb3-Dpb4, pol *ɑ*, etc., work together to ensure the balanced segregation of parental histones to both the leading and lagging strands (Gan et al. 2018; Yu et al. 2018; Petryk et al. 2018; Li et al. 2020). Our current NCOP assay does not provide information on nucleosome deposition on the leading and the lagging strands; however, the transient gaps in nucleosome occupancy behind the replication fork in the absence of *Cac*1 may affect how parental histone marks are copied and restored on the daughter chromatids.

Underscoring the importance of nucleosome deposition and maturation, loss of CAF-1 activity results in loss of gene silencing (Enomoto and Berman 1998; Monson et al. 1997; Smith et al. 1999) as well as increased cryptic transcription (van Bakel et al. 2013). Consistent with the transient delay we observed in nucleosome deposition and maturation in the absence of CAF-1, we found that the loss of silencing and cryptic transcription phenotypes were transient and confined to S-phase. Similarly, the de-regulation of the DNA replication timing program leads to S-phase-specific transcriptional changes which mimic those caused by defects in chromatin assembly factors, presumably because the chromatin assembly machinery cannot keep pace with the rapid replication rate resulting in transient defects in histone assembly (Santos et al. 2022). The S-phase-specific increase in gene expression from the subtelomeric regions of the chromosomes suggests that the re-establishment of silencing following passage of the fork was initially impaired but later recovered following chromatin maturation. We propose that the gaps in nucleosome distribution prevent the spreading of the silencing near the telomeres, likely by disrupting the nucleosome-Sir3 interaction (Brothers and Rine 2022; Saxton and Rine 2020). The elevated cryptic transcription during S-phase likely results from the promiscuous initiation of transcription at sequences associated with slow-to-mature nucleosomes. Together, these results underscore the importance of DNA replication and chromatin maturation in shaping the chromatin landscape and open up new avenues for understanding how aberrations in these processes might contribute to dynamic gene expression programs during development and in disease states such as cancer.

## Materials and Methods

### Yeast strains

All yeast strains are in the W303 background. WT cells have the genotype: MATa, leu2-3,112, BAR1::TRP, can1-100, URA3::BrdU-Inc, ade2-1, his3-11,15. *cac1Δ* cells have the genotype: MATa, leu2-3,112, BAR1::TRP, can1-100, URA3::BrdU-Inc, ade2-1, his3-11,15, rlf2*Δ*::HIS.

### Cell growth and culturing

To profile nascent and mature chromatin, yeast cells were grown at 25°C to an OD_600_ of ∼0.7. EdU (Berry & Associates) was added to the culture at a final concentration of 130 µM. At the same time, *ɑ*-factor (GenWay) was added to the culture at a final concentration of 50 ng/mL. After 5 minutes of EdU pulse, a sample was taken for nascent chromatin, after which cells were washed and resuspended in fresh media containing 130 µM thymidine (Sigma-Aldrich) and 50 ng/mL *ɑ*-factor. Samples were taken at 10, 15, 20, and 40 minutes after the media change. Cells were washed twice with sterile water before being pelleted and flash-frozen. Cell pellets were stored at −80°C. All experiments were performed with independent biological replicates.

To collect samples for RNA-seq, yeast cells were grown at 25°C to an OD_600_ of ∼0.7 and arrested in G1 phase with *ɑ*-factor at a final concentration of 50 ng/mL for 2 h. At the end of *ɑ*-factor arrest, a sample was collected (af sample), and then the cells were washed twice in sterile water and released into fresh media. Samples were collected at 10-minute intervals for 60 minutes, after which *ɑ*-factor was added back to the media at the same concentration. A final sample was collected at 150 minutes after release from the initial *ɑ*-factor. The cells were processed and stored as mentioned above. At each time point, a separate sample was collected for flow cytometry. An independent biological replicate was performed with time points af, 40 minutes, and 60 minutes. Figure 5B-E were generated with replicate 1. A comparison between the two replicates is shown in Supplemental Figure 10.

### Chromatin preparation

MNase digestions were performed as previously described (Belsky et al. 2015).

### Click reaction and streptavidin affinity capture

Sixty micrograms of MNase-digested DNA was mixed with click chemistry reaction buffers as previously described (Stewart-Morgan et al. 2019) in siliconized tubes (Bio Plas). The mixture was incubated in the dark at room temperature for 30 minutes. DNA was purified using Probe Quant G50 columns (GE Healthcare), followed by ethanol precipitation. To capture biotin-conjugated DNA, 10 µL of streptavidin magnetic beads (New England Biolabs) per sample were blocked as previously described (Gutiérrez et al. 2019). After blocking, beads were washed with cold binding buffer (1 M Tris, 5 M NaCl, 10% NP-40, 0.5 M EDTA) and resuspended in the same volume of the binding buffer as the original volume of beads. Streptavidin beads were mixed with DNA and incubated with 200 µL of binding buffer overnight at 4°C with gentle rotation. Following incubation, bead-bound DNA was washed twice with cold binding buffer and three times with EB buffer (Qiagen).

### Sequencing library preparation

Library preparations were performed on bead-bound DNA as previously described (Gutiérrez et al. 2019).

### Data analysis

Data analysis was performed in R version 4.3.0 (https://www.r-project.org). See supplemental materials for detailed descriptions of data analysis. Data processing scripts are available at https://gitlab.oit.duke.edu/bc202/chen_2023_paper

### Data Access

All sequencing data from this study are available in the NCBI Sequence Read Archive (SRA) under accession number PRJNA974469.

## Acknowledgments

We thank former and current members of the MacAlpine group and the Hartemink group for critical comments and suggestions during the development of the project. We thank Dr. Lee Zou for a critical reading of the manuscript. D.M.M. is supported by the National Institutes of Health grant R35-GM127062, and A.J.H. is supported by grant R35-GM141795. Figure 1A was created with BioRender.com.

## Author contributions

D.M.M. and B.C. designed the experiments and wrote the manuscript. B.C. performed experiments and data analysis. H.K.M. performed experiments. A.J.H. advised on data analysis and edited the manuscript.

