## Supplemental Materials for "Spatiotemporal kinetics of CAF-1-dependent chromatin maturation ensures transcription fidelity during S-phase"

### Supplemental Methods

*Sequence alignment*

All reads were aligned to the sacCer3/R61 version of the *S. cerevisiae* genome using Bowtie 0.12.7 (Langmead et al. 2009). Reads from nascent chromatin occupancy profiling were aligned in paired-end mode using the following Bowtie parameters: -n 2 -l 20 -m 1. RNA-seq reads were aligned in single-end mode with the following Bowtie parameters: -n 2 -l 20 -M 1.

*Analysis of chromatin organization*

For NCOP experiment analyses, biological replicates were merged and sampled to the same read depth for normalization. Autocorrelation function (ACF) analysis was performed to assess nucleosome organization for the first four nucleosomes (700 bp) in gene bodies downstream of the TSS. 4214 genes that have an open reading frame (ORF) longer than 700 bp and have an annotated TSS coordinate (Park et al. 2014) were selected for the ACF analysis. To calculate the ACF value, midpoint locations of nucleosome-sized reads between 140 bp and 180 bp in length were used to generate density estimates with a 30 bp bandwidth Gaussian kernel. The density estimates were used to determine the ACF value corresponding to the nucleosome periodicity lag of 172 bp (Gutiérrez et al. 2019) for each gene. ACF values from the mature chromatin of WT cells were used to select the top 50% of genes with regularly phased arrays of nucleosomes (Figure 1C).

*Analysis of nucleosome occupancy and assembly rate*

Nucleosome dyad positions were determined from the WT chase 40 minute sample, which were used as a reference for all downstream analyses. To identify the nucleosome dyad positions, nucleosome-sized reads were used to generate a density curve across the length of each chromosome with a 30 bp bandwidth Gaussian kernel. The density curve calculated from each chromosome was passed through the turnpoints function in R to determine peaks of the density curve, which were used as nucleosome dyad positions.

Nucleosome occupancy was calculated as the number of nucleosome-sized reads mapped to the 140 bp window centered around each nucleosome dyad. High-confidence nucleosomes were defined as nucleosomes with occupancy greater than the 25th percentile at the final time point in both WT and *cac1Δ* cells.

Assembly timing index (ATI) is used to assess the rate of maturation for individual nucleosomes. ATI is calculated as the inverse of the mean weighted sum of nucleosome occupancy at each time point ($O_n$) (time points 1, 2, 3, 4, 5 represent pulse 5', chase 10', 15', 20', 40'):

$$ATI = 1/\frac{\sum_{n=1}^{5} n * O_n}{5}$$

*Fragile nucleosomes*

To determine the positions of fragile nucleosomes, we used MNase-seq data generated with MNase concentrations of 200 U and 400 U (Chereji et al. 2017), and identified nucleosomes that existed in the 200 U sample but not in the 400 U sample. To do so, we first used RoboCOP (Mitra et al. 2021) to call nucleosomes separately in the 200 U sample and 400 U sample. Then, using the match_pair_nucs() function in RoboCOP, the nucleosomes called from the two samples were compared as follows: 1) If prob(nuc200)>= 0.1 and prob(nuc400) >=0.1 and the two dyads were less than 73 bp apart, then the two nucleosomes were considered to be the same nucleosome. 2) If prob(nuc200) >=0.1 or prob(nuc400)>=0.1 and no other nucleosomes in the other data set with probability >=0.1 were within 73 bp, the nucleosome is considered to be unique to the sample.

*Genomic features of slow and fast nucleosomes*

A single intergenic region is defined as the region between two neighboring nonoverlapping ORFs. A nucleosome is considered to be located in a gene if the nucleosome dyad lies within the ORF of the gene.

To compute the dinucleotide frequency for the fast and slow nucleosomes, we first matched the dyad positions of fast and slow nucleosomes identified from this study to nucleosome positions determined from chemical mapping (Brogaard et al. 2012) by finding their nearest counterparts. To be considered a match, the two dyads need to be less than 50 bp apart. Dinucleotide frequency was generated by aligning the matched nucleosome positions from chemical mapping to the sacCer2 reference genome.

*Analysis of subnucleosomal fragments*

An autocorrelation function was used to determine the phasing of the subnucleosomal fragments in the nascent chromatin of *cac1Δ* cells. Midpoints of subnucleosomal fragments located within the first 700 bp of each gene were used to generate ACF values corresponding to lags ranging from 20 bp to 200 bp. The phasing for subnucleosomal fragments in each gene is determined as the lag that maximizes the ACF value.

### Supplemental Figures

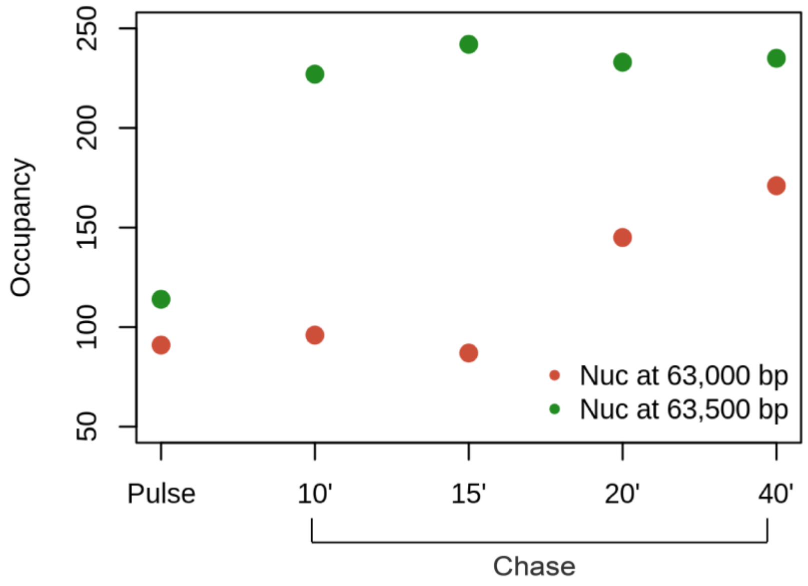


**Supplemental Figure 1.** NCOP captures nucleosomes assembled at heterogeneous rates in *cac1Δ* cells. Occupancy of nucleosomes positioned at 63,000 bp and 63,500 bp on ChrI in *cac1Δ* cells across all time points.

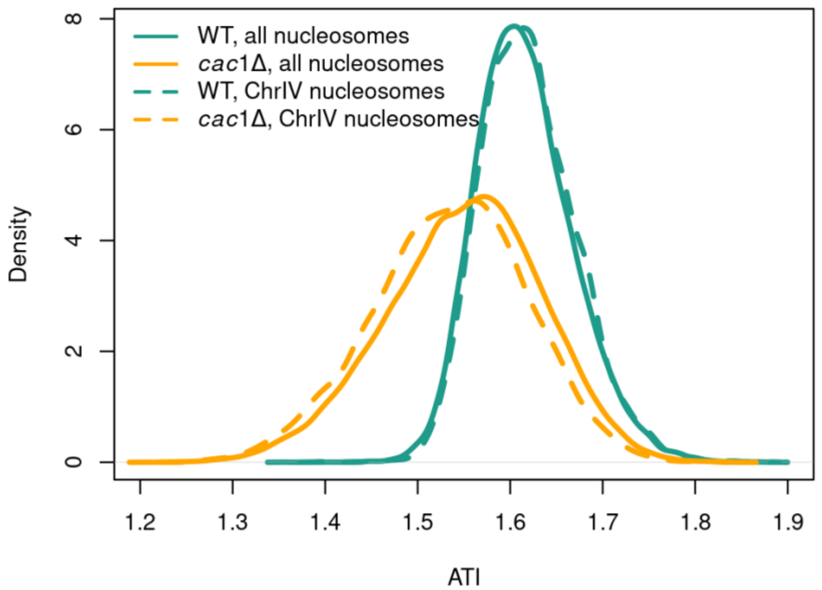


**Supplemental Figure 2.** Nucleosomes on ChrIV are representative of nucleosomes across the genome. Density distribution of ATI values for all high-confidence nucleosomes and nucleosomes on ChrIV in WT and *cac1Δ* cells.

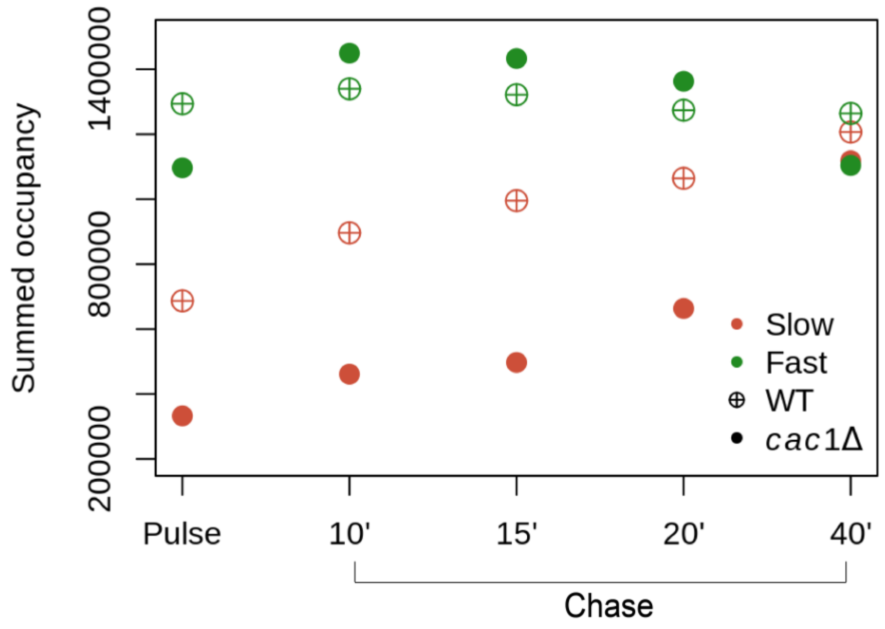


**Supplemental Figure 3.** Fast and slow nucleosomes identified in *cac1Δ* cells have different maturation kinetics. Summed occupancy of fast and slow nucleosomes identified in WT and *cac1Δ* cells across all time points.

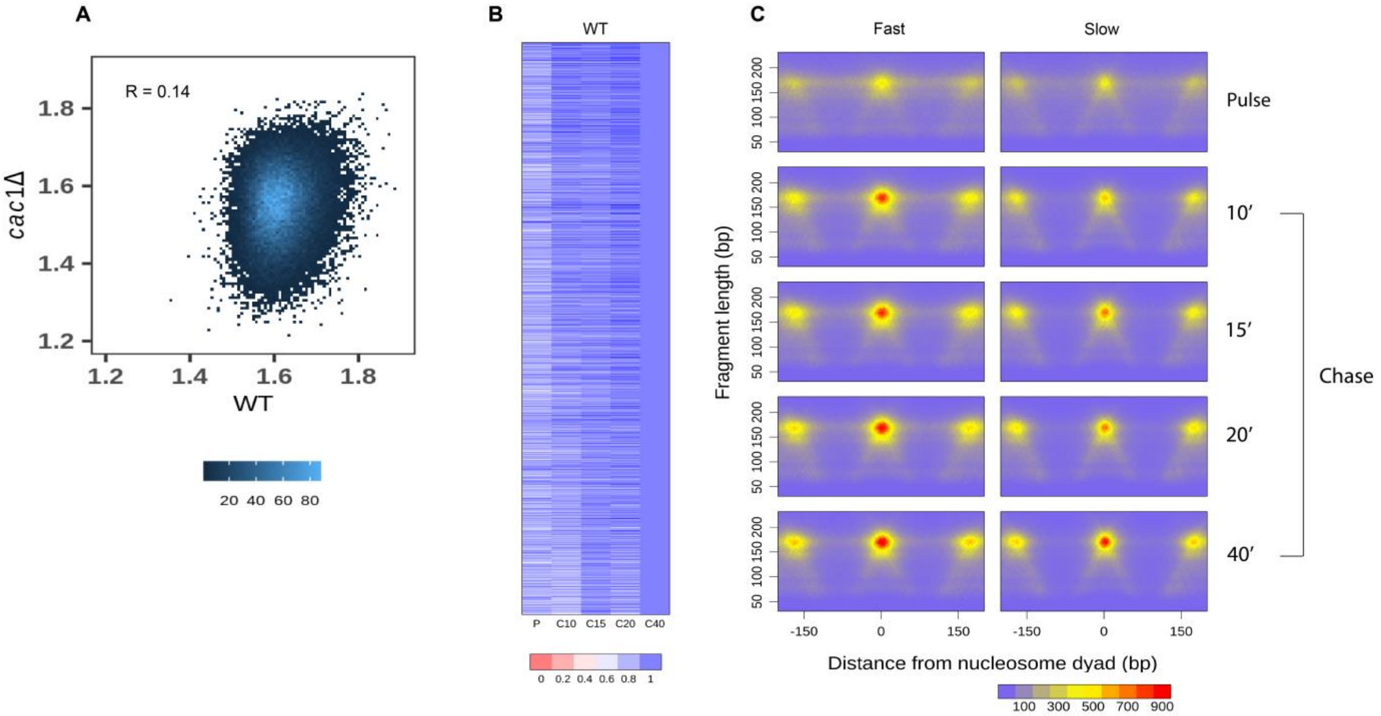


**Supplemental Figure 4.** Nucleosomal ATI values determined from WT and *cac1Δ* cells are not correlated with each other. (A) Density scatter plot depicting the ATI values for individual nucleosomes from WT and *cac1Δ* cells. (B) Heatmap showing the nucleosome occupancy at each time point relative to the final time point in WT, presented in order of decreasing ATI values determined from *cac1Δ* cells. (C) Aggregate chromatin profiles of slow and fast nucleosomes identified from *cac1Δ* cells using WT data.

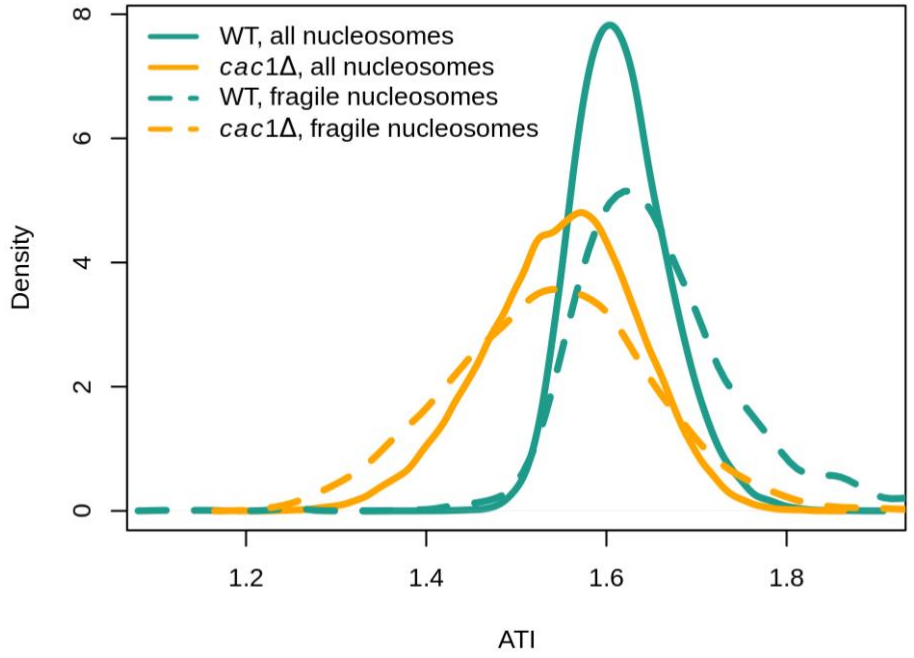


**Supplemental Figure 5.** Maturation kinetics of nucleosomes are not related to their sensitivity to MNase digestion. Density distribution of ATI values for all high-confidence nucleosomes and fragile nucleosomes in WT and *cac1Δ* cells.

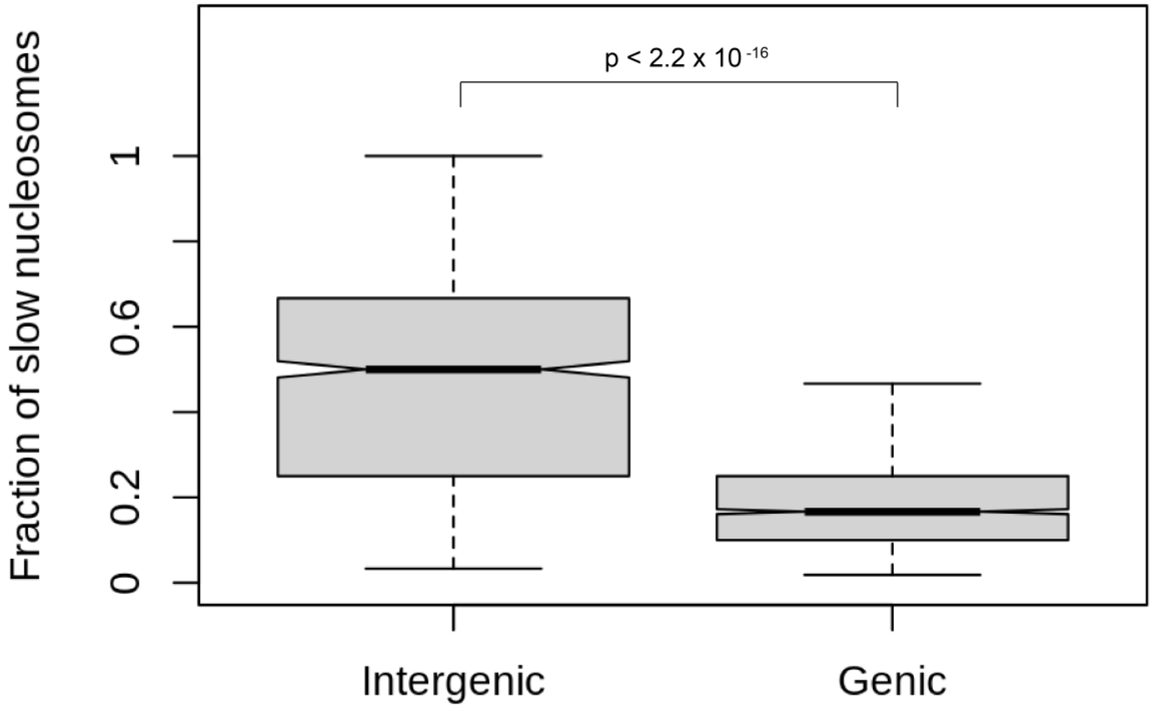


**Supplemental Figure 6.** Slow nucleosomes are enriched in intergenic regions compared to gene bodies. Boxplot depicting the fraction of slow nucleosomes, identified in *cac1Δ* cells, among all nucleosomes in the intergenic regions and genic regions. The difference between the fraction of slow nucleosomes in the intergenic regions and genic regions is significant ($p < 2.2 \times 10^{-16}$ ).

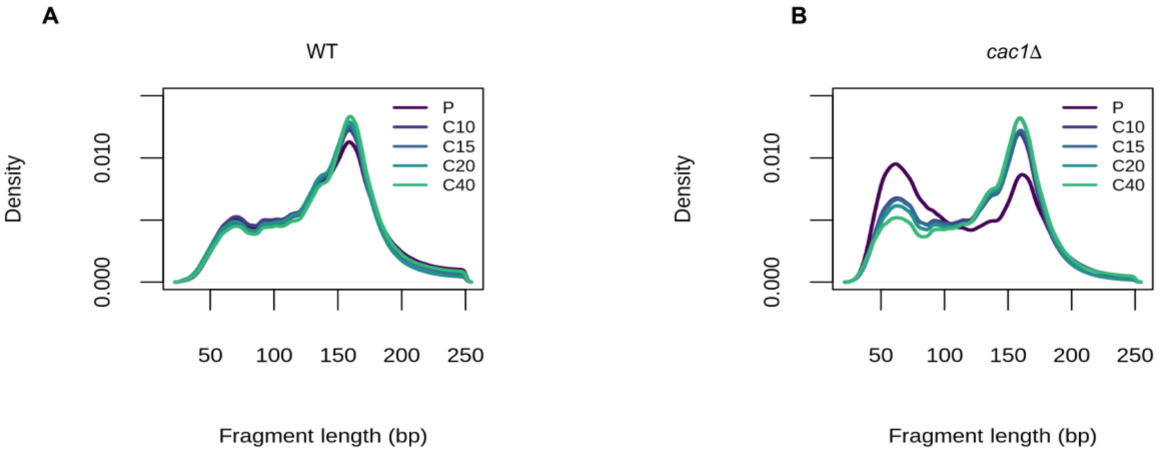


**Supplemental Figure 7.** The nascent chromatin of *cac1Δ* cells displays an increase in subnucleosomal-sized fragments that gradually disappears as the chromatin matures. Density distribution of fragment length in (A) WT and (B) *cac1Δ* cells at all time points.

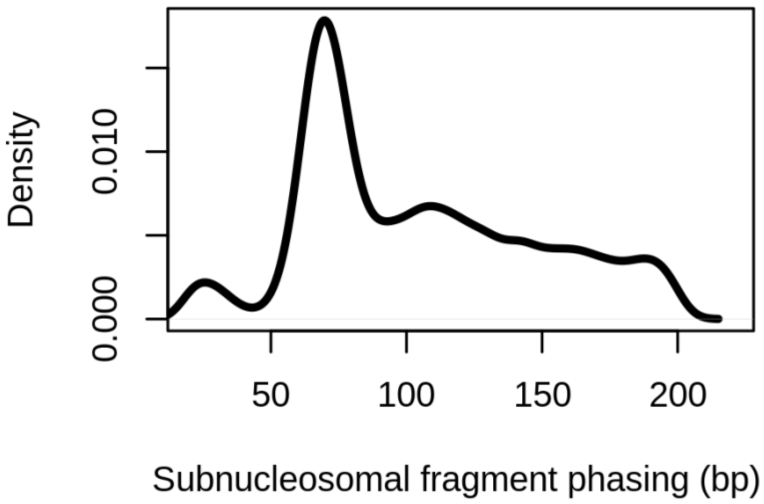


**Supplemental Figure 8.** Subnucleosomal-sized fragments captured from nascent chromatin in *cac1Δ* cells form ordered arrays of phased clusters. Density distribution of the phasing of subnucleosomal fragments in gene bodies in the nascent chromatin of *cac1Δ* cells.

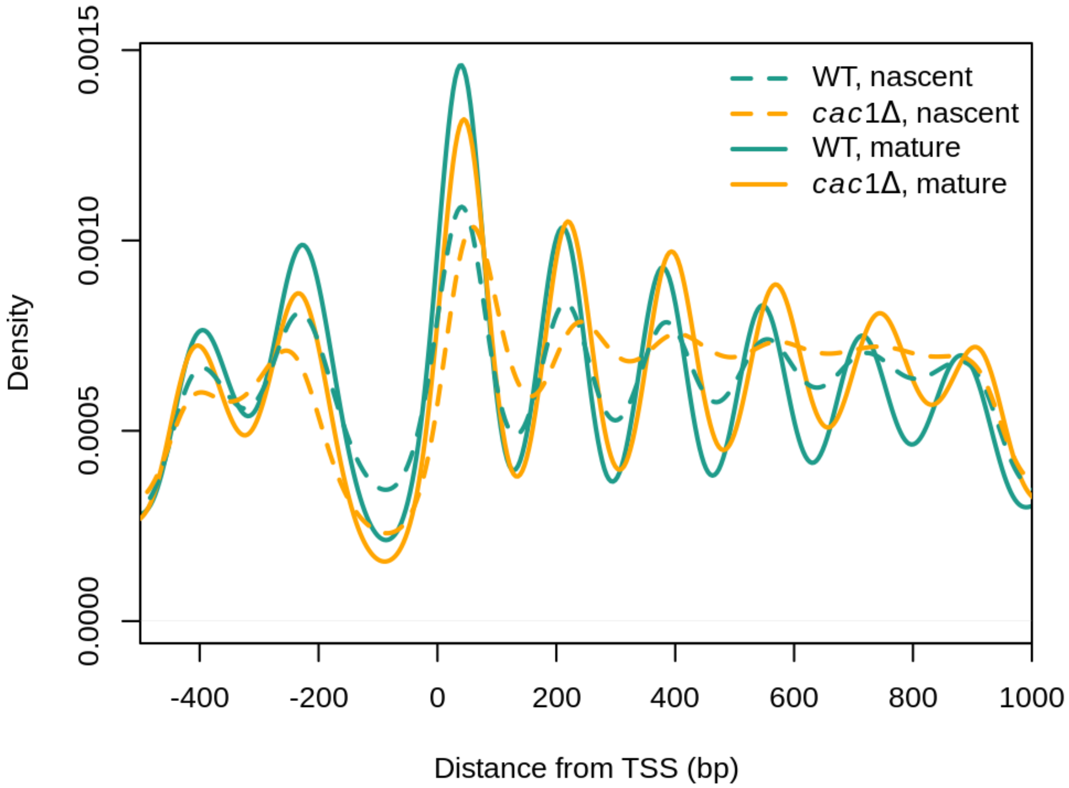


**Supplemental Figure 9**. Nucleosome occupancy profiles of WT and *cac1Δ* cells after a 5 minute EdU pulse (nascent) and 40 minutes following a thymidine chase (mature).

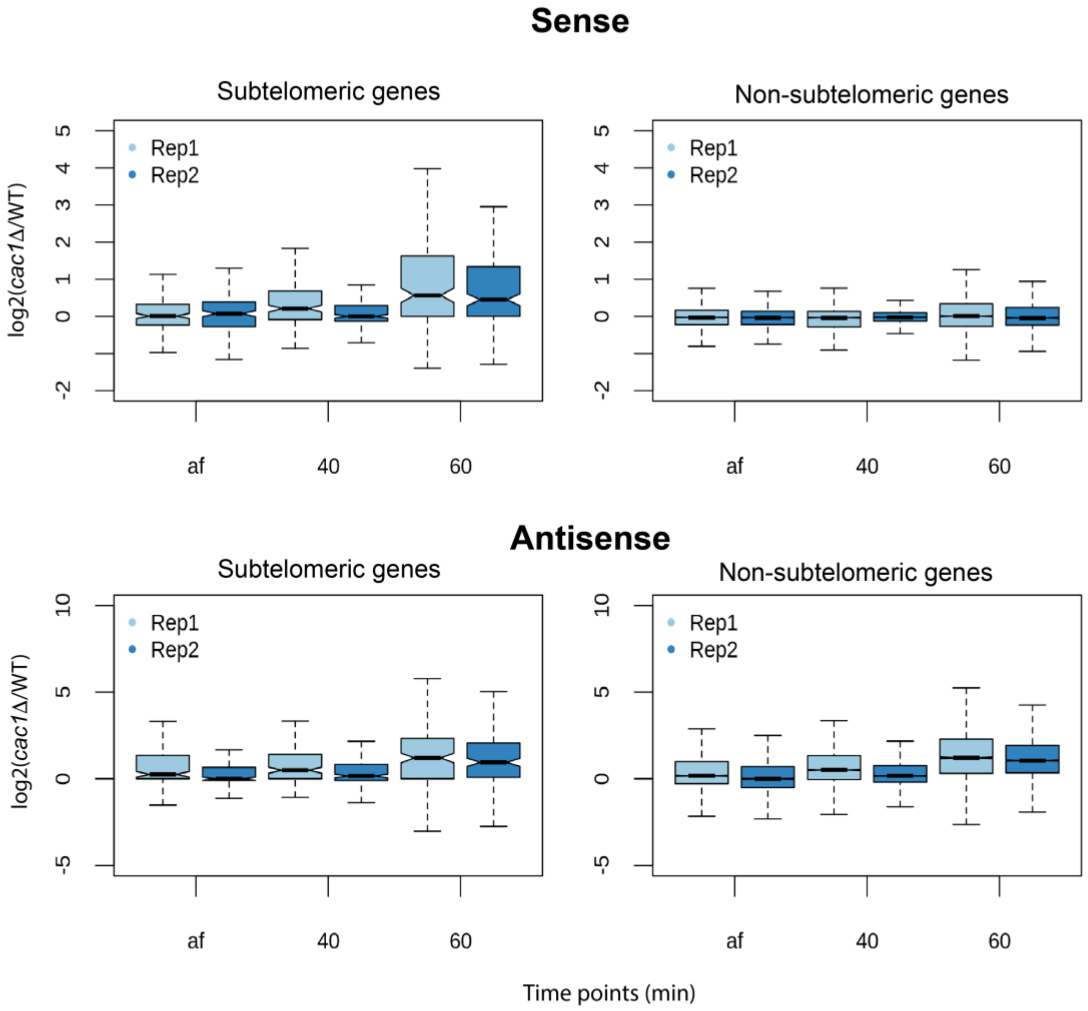


**Supplemental Figure 10.** Two biological replicates of the RNA-seq experiments are consistent with each other. These four panels show data from experiments undertaken to replicate the results of Figures 5B-E.
